# Emergence of the subapical domain is associated with the midblastula transition

**DOI:** 10.1101/713719

**Authors:** Anja Schmidt, Jörg Großhans

## Abstract

Epithelial domains and cell polarity are determined by polarity proteins which are associated with the cell cortex in a spatially restricted pattern. Early Drosophila embryos are characterized by a stereotypic dynamic and de novo formation of cortical domains. For example, the subapical domain emerges at the transition from syncytial to cellular development during the first few minutes of interphase 14. The dynamics in cortical patterning is revealed by the subapical markers Canoe/Afadin and ELMO-Sponge, which widely distributed in interphase 13 but subapically restricted in interphase 14. The factors and mechanism determining the timing for the emergence of the subapical domain have been unknown. In this study, we show, that the restricted localization of subapical markers depends on the onset of zygotic gene expression. In contrast to cell cycle remodeling, the emergence of the subapical domain does not depend on the nucleo-cytoplasmic ratio. Thus, we define cortical dynamics and specifically the emergence of the subapical domain as a feature of the midblastula transition.

**Author summary:** Midblastula transition is a paradigm of a developmental transition. Multiple processes such as cell cycle, cell mobility, onset of zygotic gene expression, degradation of maternal RNA and chromatin structure are coordinated to lead to defined changes in visible morphology. The midblastula transition in *Drosophila* embryos is associated with a change from fast nuclear cycles to a cell cycle mode with gap phase and slow replication, a strong increase in zygotic transcription and cellularization. The timing of the processes associated with the midblastula transition are controlled by the onset of zygotic gene expression or the nucleocytoplasmic ratio. Here we define the patterning of cortical domains, i. e. the emergence of a subapical domain as a novel feature of the midblastula transition whose appearance is controlled by the onset of zygotic transcription but not the nucleocytoplasmic ratio. Our findings will help to gain further understanding of the coordination of complex developmental processes during the midblastula transition.

## Introduction

The cell cortex underlies the plasma membrane and consists of a layer of F-actin and associated proteins, including actin nucleators, regulators and myosin motors. Proteins, such as ERM proteins, link F-actin to the plasma membrane (1). Typical for epithelial cells are cortical domains, which contain marker proteins specific for the respective domain in addition to the general set of cortical proteins. For example, Par-3/Bazooka (Baz) typically marks the subapical domain, whereas Par-1 marks the lateral domain (2,3). Although mutual exclusion of such marker proteins has been shown to maintain boundaries between two domains in some cells, the mechanism for initial establishment of the domains and pattern formation is not well defined. The de novo appearance of the first epithelium during cellularization in *Drosophila* embryos, provides an excellent model to study the initial formation of cortical domains and epithelial polarization (4).

Following a syncytial phase of development with rapid nuclear cycles typical for insects, the first epithelium forms after about two hours of embryonic development as a morphologically obvious feature marking the transitions from syncytial to cellular blastoderm (5–7). This morphological change, often referred to as midblastula transition (MBT) is associated with several cellular processes that appear to be coordinated, including remodeling of the cell cycle, transition to a slow mode of DNA replication, heterochromatin formation, ingression of the cellularization furrow, elongation of the nuclei, and importantly activation of the zygotic genome (6,8,9). Concerning epithelial polarization it is important to note that the number of cortical domains increases during the transition from two cortical domains (caps and intercaps) in interphase 13 (10,11) and three domains (apical, lateral, basal) during mitosis (12) to the typical four domains. A dedicated subapical region positioned between the apical and lateral domains emerges for the first time in development in interphase 14 (3,8).

It is unknown, if and how the emergence of the subapical domains is linked or coordinated with the other processes associated with the midblastula transition. It has been previously shown that zygotic transcription initiates the cell cycle remodeling and is required for cellularization (13). The changes are due to specific zygotic genes, e. g. *slam, nullo, frs* or to global signals such as transcription associated DNA replication stress and DNA checkpoint activation (13). The emergence of the subapical domain has not been investigated in this context, so far.

The earliest marker proteins for the prospective subapical domain during onset of cellularization are Canoe (Cno, Afadin in vertebrates) and the unconventional GEF complex ELMO-Sponge (8,14), which act upstream of Canoe possibly via control of the small GTPase Rap1. Both Canoe and ELMO-Sponge are widely distributed during the syncytial interphases and mitoses (nuclear cycles 10–13). Canoe is detected in cap and intercap regions, whereas the ELMO-Sponge complex marks the actin caps and control their formation (8). This disc-like pattern in pre-MBT interphases changes to a ring-like pattern in interphase 14, when ELMO-Sponge initiate restriction of Canoe to the prospective subapical region. Only during the course of cellularization, the typical subapical proteins Bazooka/Par-3 and Armadillo (Arm, β-Catenin in vertebrates) are enriched in the subapical region (15–17).

In this study, we investigate the role of zygotic gene expression and cell cycle remodeling for the formation of the subapical domain. As Bazooka feeds back on subapical restriction of Canoe later in cellularization, we tested the function of this genetic interaction for the initial emergence of the subapical region. We show that the localization of early subapical domain markers like ELMO-Sponge and Canoe depends on onset of zygotic expression but not cell cycle remodeling and not on *bazooka* during early cellularization.

## Results

### Change of Canoe distribution pattern at the onset of interphase 14

The subapical cortical domain emerges during the transition from syncytial to cellular blastoderm for the first time during embryonic development. During this process the localization pattern of the actin binding protein Canoe changes from a dispersed pattern at the actin caps to a coalesced pattern at the prospective subapical domain within about five minutes of the onset of cellularization in interphase 14 (Figure 1A) (8). Subapical restriction of Canoe depends on the small GTPase Rap1 and the unconventional guanyl nucleotide complex ELMO-Sponge, which undergoes a relocalization from discs in interphase 13 to rings in interphase 14 (8). New cellularization furrows form between the daughter nuclei. After reached longest extension in metaphase, these furrows gradually retract in the second half of mitosis to a length of about 3 µm (32,33) (Figure 1B). We applied our live imaging assay with embryos expressing the subapical marker CanoeYFP and basal marker CherrySlam to reveal the kinetics of marker segregation. Axial stacks were recorded and computationally projected to sagittal sections. During mitosis, Cherry Slam was detected at the tip of the metaphase furrow, whereas CanoeYFP was spread along the full length (Figure 1C). It is important to note the difference between “old” cellularization furrows, which arise from retracting metaphase furrows, and “new” cellularization furrows, which ingress between daughter nuclei. In “new” cellularization furrows CanoeYFP associates within minutes to the in folding membrane. In contrast, Canoe distribution is becomes subapically restricted at “old” furrows starting from a wide distribution along the furrow (Figure 1C). CherrySlam remains at the tip of “old” furrows, and gradually appears at the tip of “new” furrows (Figure 1C) (8). Although we and others have uncovered the mechanism for subapical restriction of Canoe (3,8,15,17,34), the factors determining the timing have not be studied.

**Figure 1.**
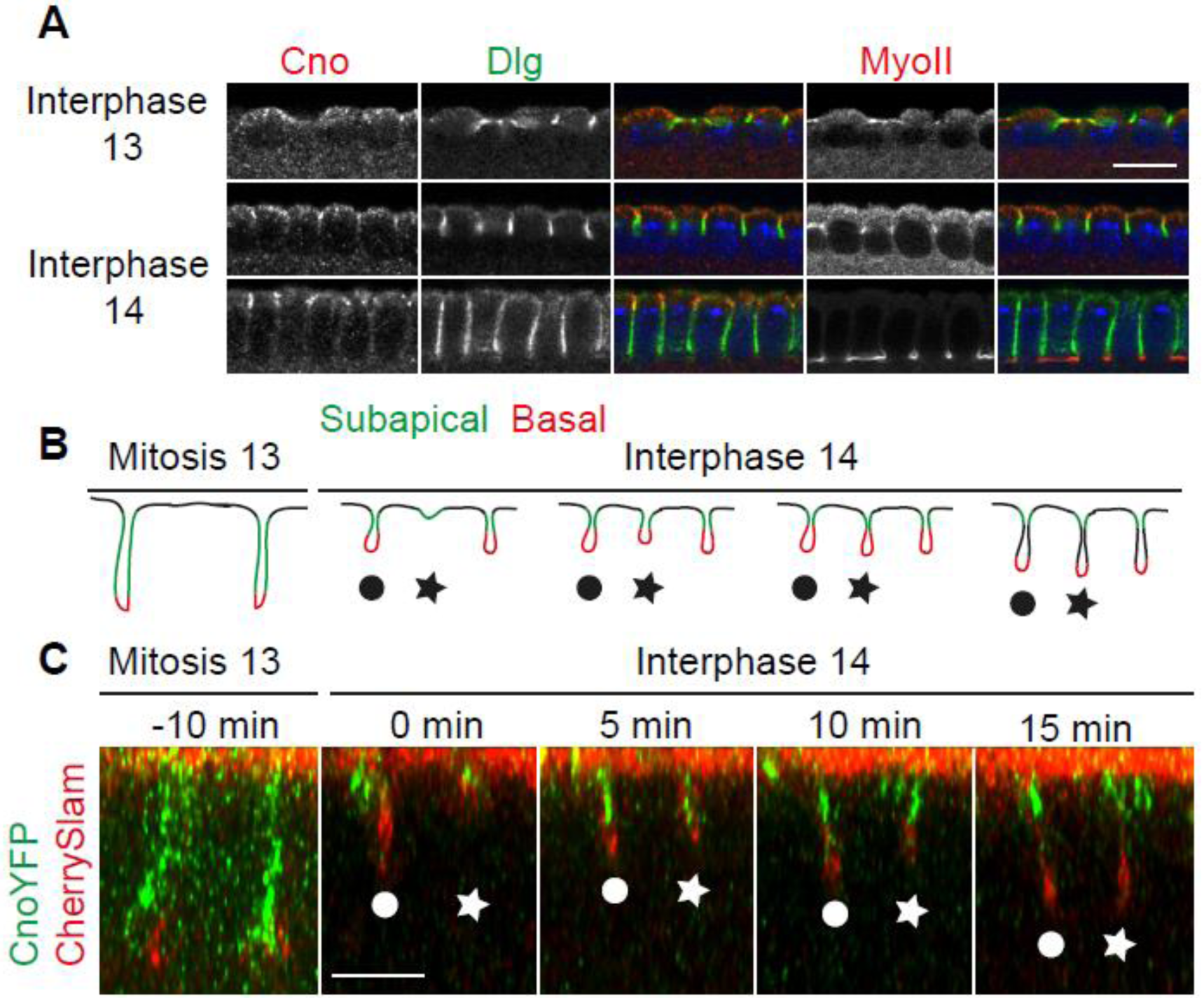
Canoe relocalizes in old cellularizatiation furrows arising from retracting metaphase furrows. (A) Fixed wild type embryos during interphases 13 and 14 as indicated stained against Canoe (grey/red), Dlg (grey/ green), MyosinII (grey/ red) and DNA (blue). (B) Scheme of metaphase and cellularization furrows during mitosis 13 and switch to interphase 14 as indicated. Proteins localizing to the subapical domain during interphase 14 (green) localize to the whole cortex of metaphase furrows. After mitosis 13 metaphase furrows retract and come to a halt forming “old furrows” (circle) while “new furrows” form between (star). The furrwos move inwards synchronosly when they have reached the same length. (C) Living embryos expressing CanoeYFP (green) and CherrySlam (red) to mark subapical and basal domains. Orthogonal views are shown. Stages are as indicated. Scale bar 10 µm.

### Formation of the subapical domain depends on zygotic gene expression but not the nucleo-cytoplasmic ratio

The change from syncytial to cellular blastoderm at the onset of cell cycle 14 and cellularization requires zygotic gene expression (35). Although it is clear that cellularization depends on zygotic gene expression, its functional relationship to the emergence of the subapical domain has not been investigated. ELMO, Sponge, Rap1 and Canoe are maternally derived proteins, whose total levels are assumed not to change much. Rather, their distribution on the plasma membrane is controlled by post-translational mechanisms.

We first asked whether the spatial restriction of subapical markers depended on zygotic transcription. We analyzed embryos, in which zygotic transcription was blocked by α-amanitin, an efficient inhibitor of RNA polymerase II. Injection of α-amanitin impairs furrow invagination and cellularization (35). Early embryos expressing CanoeYFP were injected with α-amanitin prior to cellularization (Figure 2A). In fixed control embryos, Sponge and CanoeYFP marked the invaginating furrows in a hexagonal pattern, enclosing the nuclei as visible in surface views (Figure 2A). In contrast, no spatial restriction of Sponge and CanoeYFP was detected in injected embryos in interphase 14, indicating that the restriction of the subapical markers depends on zygotic transcription (Figure 2B). To better resolve the dynamics and staging of the embryos, we recorded time lapse images of embryos expressing ELMO-GFP (Figure 2C-D). Control embryos showed a stereotypic ELMO localization at caps during syncytial blastoderm stages and transition to subapical rings during the first few minutes of cellularization in interphase 14 (Figure 2C). In embryos treated with α-amanitin, the cap staining during syncytial blastoderm stage was comparable to control embryos. In contrast, the ELMO-GFP signal remained widely distributed over the whole cortex without any obvious spatial restriction after mitosis 13 (Figure 2D). This loss of restriction of ELMO-GFP was observed at a time when the morphologically visible furrows has not yet formed in control embryos. Our data indicate that spatial restriction of ELMO, Sponge and Canoe in interphase 14 and thus formation of the subapical domain depends on zygotic transcription.

**Figure 2.**
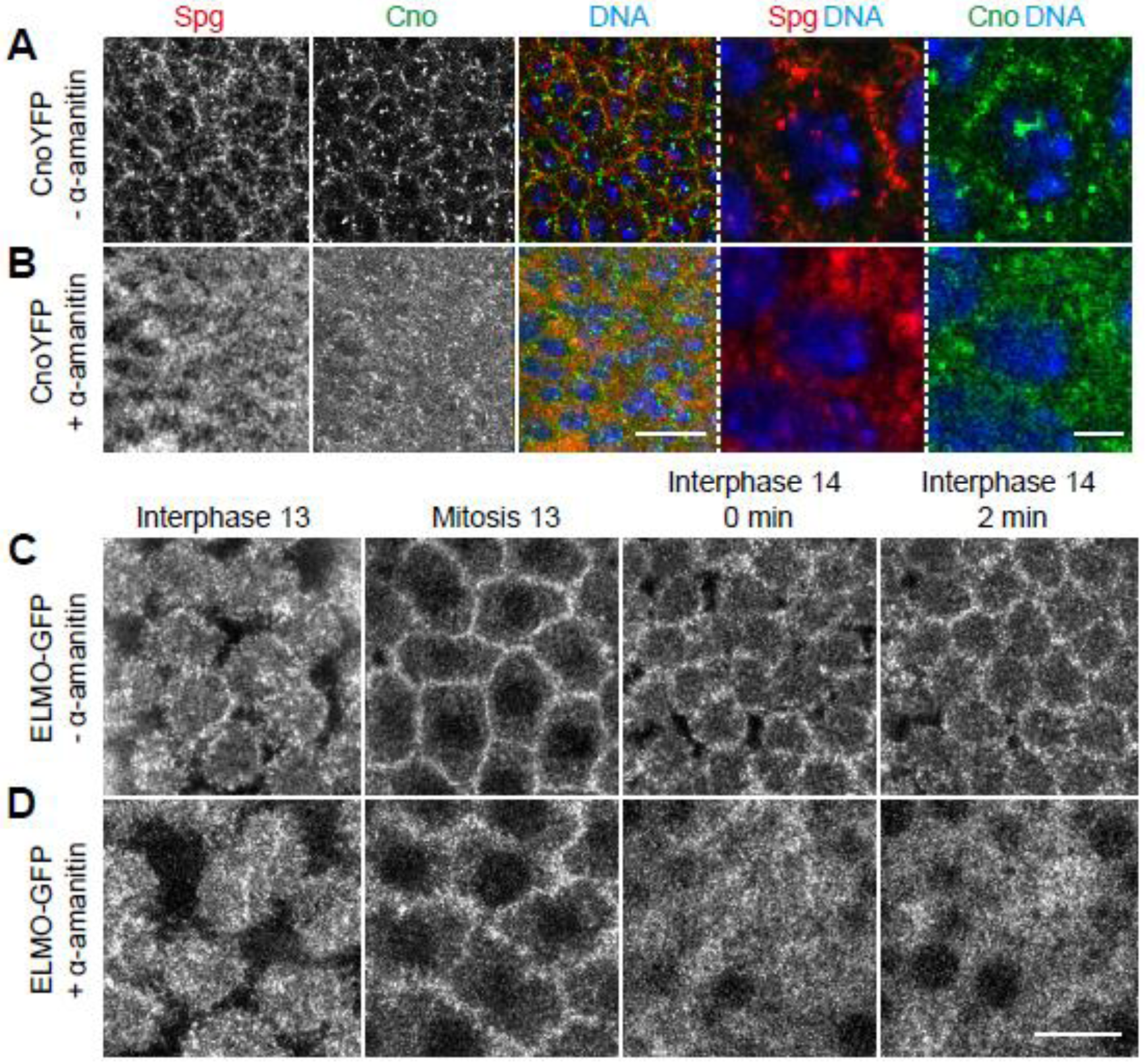
Zygotic gene expression is necessary for the formation of the subapical domain during cellularization. (A, B) Fixed non-injected (A) and α-amanitin-injected (B) embryos expressing CanoeYFP stained against Sponge (grey/ red), CanoeYFP (grey/ green) and DNA (grey/ blue) during interphase 14. Merged images and zoom-ins are shown in right panels. (B, D) Top views of images from time lapse movies of non-injected (C) and α-amanitin-injected (D) embryos expressing ELMO-GFP. Time points are indicated above the images. Scale bars 10 µm.

A second obvious timing mechanism beside onset of zygotic transcription during the transition from syncytial to cellular blastoderm is the nucleocytoplasmic ratio. Haploid embryos undergo an extra nuclear division and cellularize only in interphase 15 (36) (Figure 3C, D). It was previously reported that haploid embryos showed features of a cellularization furrow already in interphase 14, i. e. transient basal accumulation of Myo II at the furrow tip (37,38). Cortical domains have otherwise not been specifically investigated in haploid embryos, yet. We fixed and stained haploid embryos from *sésame* (*ssm, Hira*) females (21) for Canoe and F-actin. We detected specific subapical restriction of Canoe in cellularizing embryos in interphase 14 as well as in interphase 15 (Figure 3 C, D). Consistent with the previously reported basal restriction of MyoII, these data suggest that the transient furrow during interphase 14 in haploid embryos contains a patterned cortex with a subapical region. We conclude that the emergence of the subapical and basal domains does not depend on the nucleocytoplasmic ratio.

**Figure 3.**
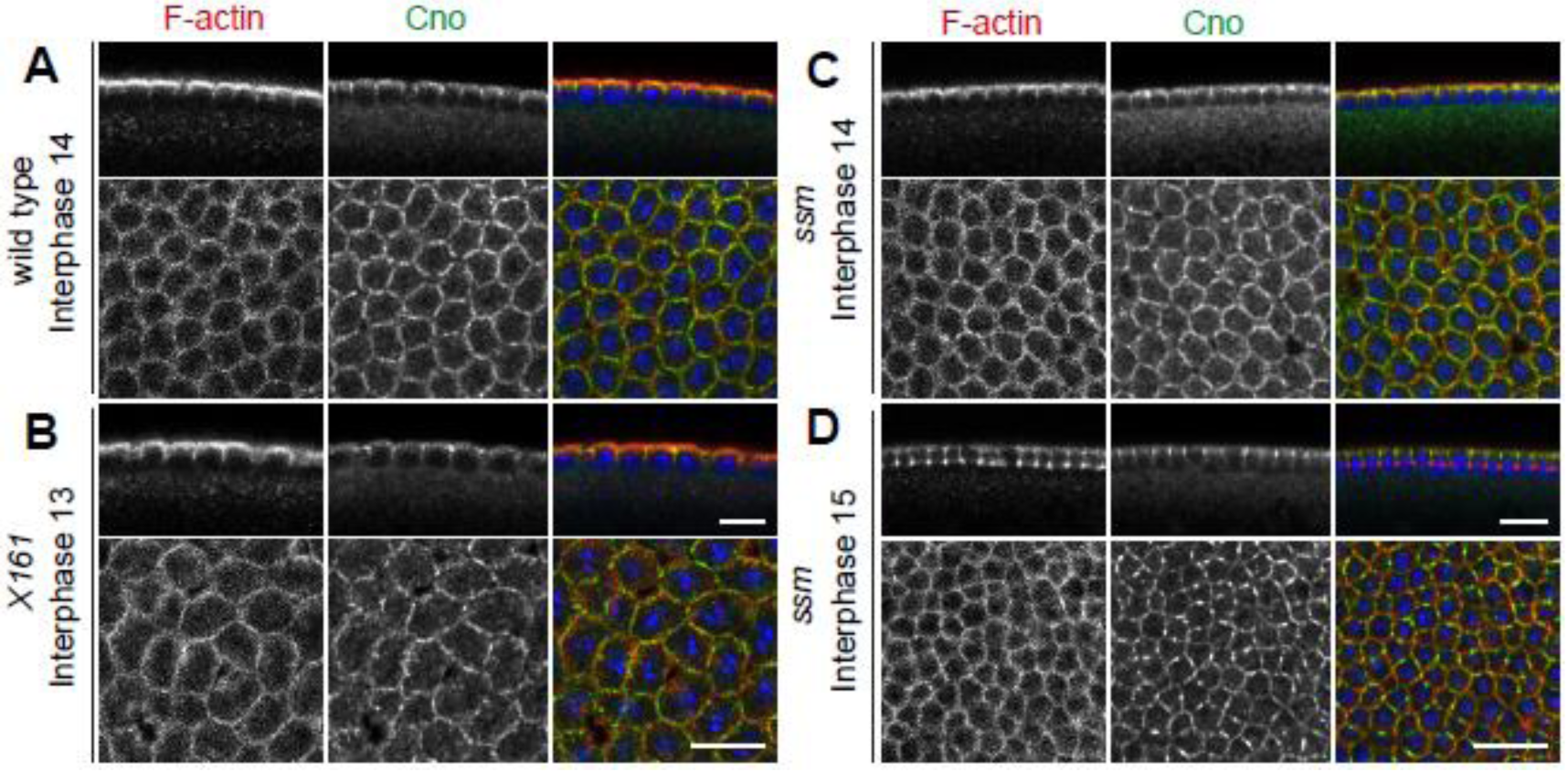
Cortical domain formation depends on zygotic gene expression and not on nucleocytoplasmic ratio. (A-D) Fixed wild type (A), *X161* (B) and *sésame* (C, D) embryos stained against F-actin (grey/ red), Canoe (grey/ green) and DNA (blue). Merged images are shown in right panels, sagittal sections in top panels and accompanying top views in lower panels. Stages are as indicated. Scale bars 10 µm.

A third timer associated with the midblastula transition is the remodeling of the fast nuclear cycle to a slow cell cycle, which depends on the onset of zygotic transcription (7). We tested whether subapical Canoe restriction would respond to a precocious zygotic transcription and precocious cell cycle remodeling. We analyzed embryos from *RPII215*^*X161*^ germline clones, which precociously start zygotic transcription, cellularize already in interphase 13, and further develop with half of the number of nuclei (9). By staining of fixed embryos, we detected a normal pattern of F-actin and subapical restriction of Canoe in embryos cellularizing in interphase 13 (Figure 3A, B).

In summary, our data indicate that the formation of the subapical domain is a regulated feature of the midblastula transition, which responds to zygotic gene expression but not on the nucleocytoplasmic ratio. The failed spatial restriction in embryos with impaired zygotic transcription may be due to the absence of one or multiple specific zygotic factors, which control the distribution pattern of ELMO-Sponge complex, for example. Alternatively, failed spatial restriction may be a consequence of zygotic transcription, such as high polymerase activity or transcription dependent DNA replication stress. Although our time lapse analysis of ELMO-GFP and CanoeYFP indicates that subapical restriction precedes furrow ingression, we do not exclude the possibility that subapical restriction is a consequence of furrow formation due to the limited resolution of our assay.

### Bazooka does not regulate subapical Canoe localization during early cellularization

Bazooka is a potential zygotic factor controlling subapical restriction of ELMO-Sponge and Canoe, since Bazooka has a maternal and zygotic expression. Previous work revealed a positive feedback mechanism during late cellularization in which subapical restriction of Canoe becomes partially dependent on *bazooka* (15). We asked whether this feedback interaction was active also during the onset of cellularization. Firstly, we analyzed the distribution of Bazooka and Armadillo which marks E-Cadherin junctions in fixed wild type embryos. For this overview, we imaged all embryos with the same laser settings to compare protein localization and amounts in different stages. With these settings, we did not detect Bazooka at Armadillo positive metaphase furrows during mitosis 13 (Figure 4A). The subapical restriction of Armadillo matures during the course of cellularization starting from an initially wide distribution along the furrow. The basal junction, in comparison, was detected very early on as reported previously (3,39)(Figure 4B–D). A clear subapical Bazooka restriction was first detected during cellularization when the furrows extended to around half the length of the elongated nuclei (Figure 4D). Remarkably, at this time point subapical Armadillo enrichment was not visible yet. During the course of cellularization Bazooka puncta persisted at the subapical position colocalizing with Armadillo (3,40) (Figure 4E, F). The lack of a subapical Bazooka signal during early cellularization does not support an early function of the feedback regulation on Canoe.

**Figure 4.**
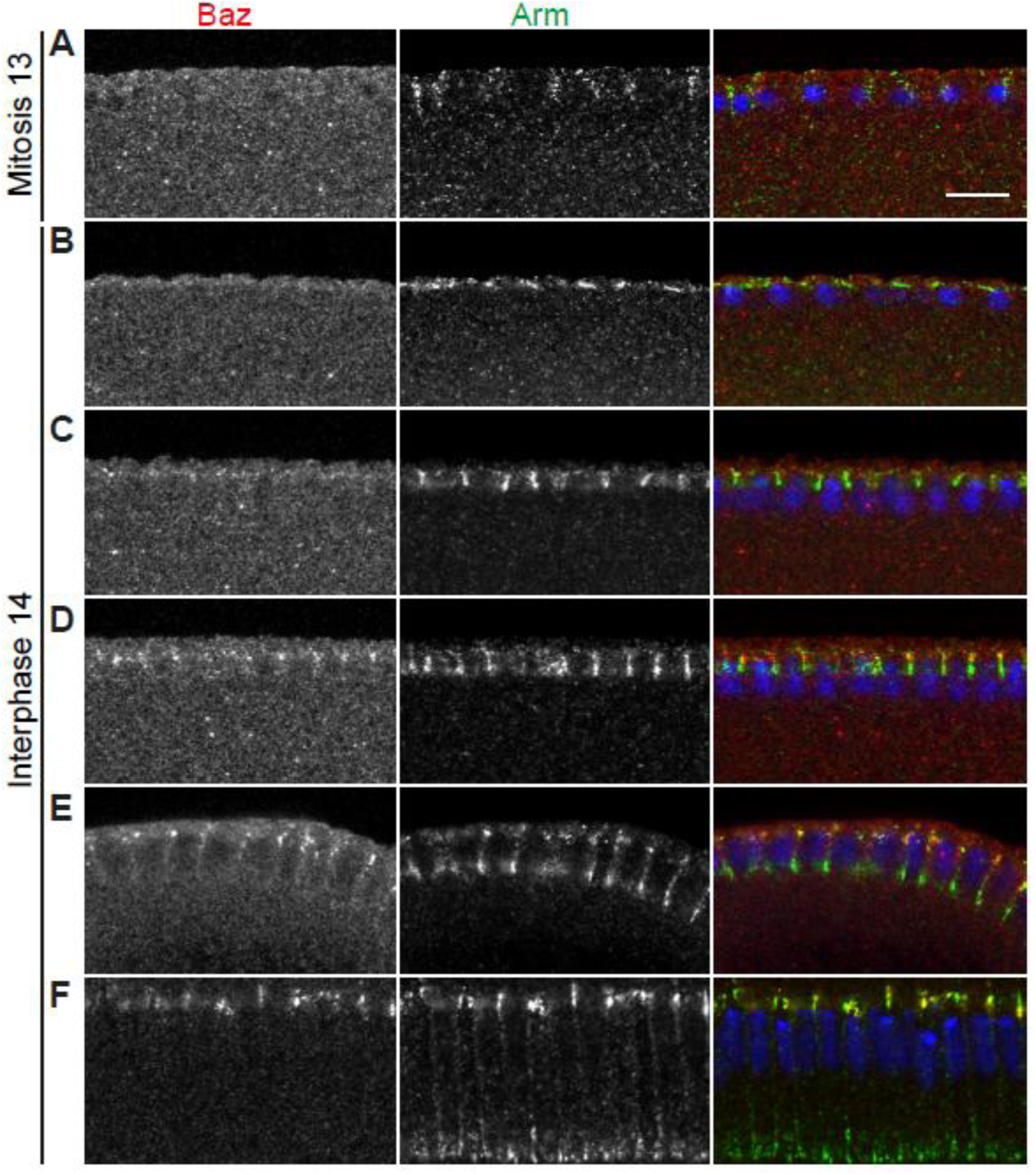
Subapical enrichment of Bazooka and Armadillo during cellularization. (A-F) Fixed wild type embryos during mitosis 13 (A) and interphase 14 (B-F, from early to late cellularization). Embryos were stained in the same tube against Bazooka (grey/red), Armadillo (grey/ green) and DNA (blue) and imaged with same laser settings to estimate different protein amounts in different stages. Scale bar 10 µm.

To clarify the relation of Canoe and Bazooka in functional terms, we depleted *bazooka* by RNAi and analyzed fixed and stained embryos (Figure 5). RNAi depletion is functional as indicated by the loss of Bazooka staining and the later phenotype with holes in the amnioserosa (Supplemental figure 1). Subapical Canoe enrichment was comparable in wild type controls and *bazookaRNAi* embryos during early cellularization whereas Canoe localization was affected as described before in early gastrulating embryos (15) (Supplemental figure 1B). Based on these data we conclude that the Bazooka-Canoe feedback loop becomes activated only during the course of cellularization and is not involved in the initial subapical restriction of Canoe.

**Figure 5.**
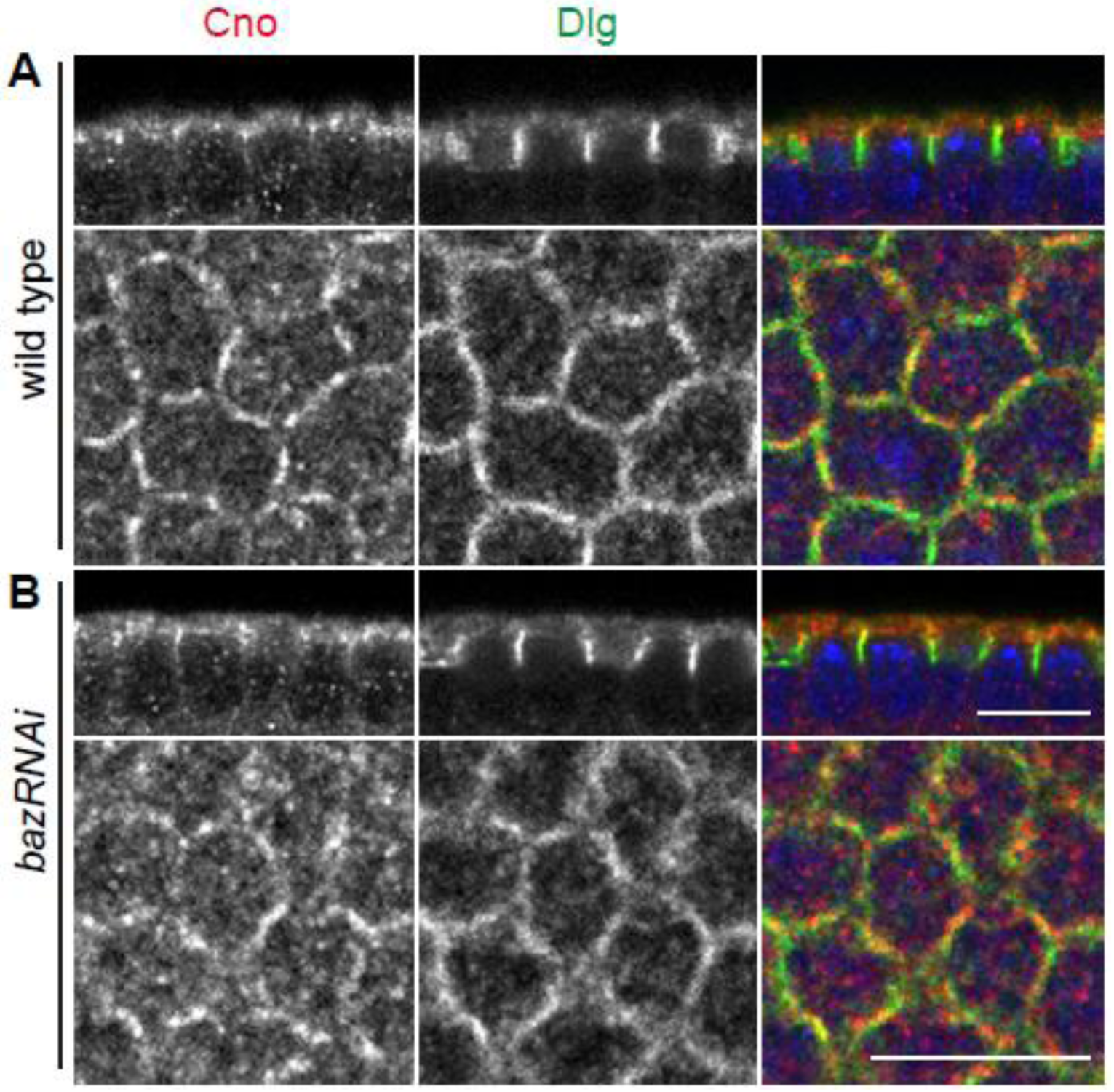
Canoe localization is not affected by Bazooka during early cellularization. (A-B) Fixed wild type (A) and *bazRNAi* (B) embryos during early cellularization stained against Canoe (grey/ red), Dlg (grey/ green) and DNA (blue). Side views are shown in upper panels and corresponding top views in lower panels. Scale bars 10 µm.

### Subapical Bazooka enrichment is controlled by the unconventional GEF ELMO-Sponge

Initial subapical restriction of Canoe is controlled by the unconventional GEF complex ELMO-Sponge and the GTPase Rap1. We asked whether this functional dependence also holds true for Bazooka and Armadillo. By analysis of fixed embryos, we found that both *Rap1* and *ELMO* were required for subapical restriction of both Bazooka and Armadillo. Bazooka and Armadillo staining was dispersed along the lateral furrow in embryos from females with ELMO as well as Rap1 germline clones consistent with previous reports (15) (Figure 6B, D). Conversely Bazooka and Armadillo did not depend on a different Rap1GEF, *dizzy*, (Figure 6C) consistent with our previous report that subapical restriction of Canoe did not depend on *dizzy* (8). These findings confirm the earlier described pathway of Bazooka being downstream of the unconventional Rap1 GEF complex ELMO-Sponge, Rap1GTPase and Canoe during early and late cellularization.

**Figure 6.**
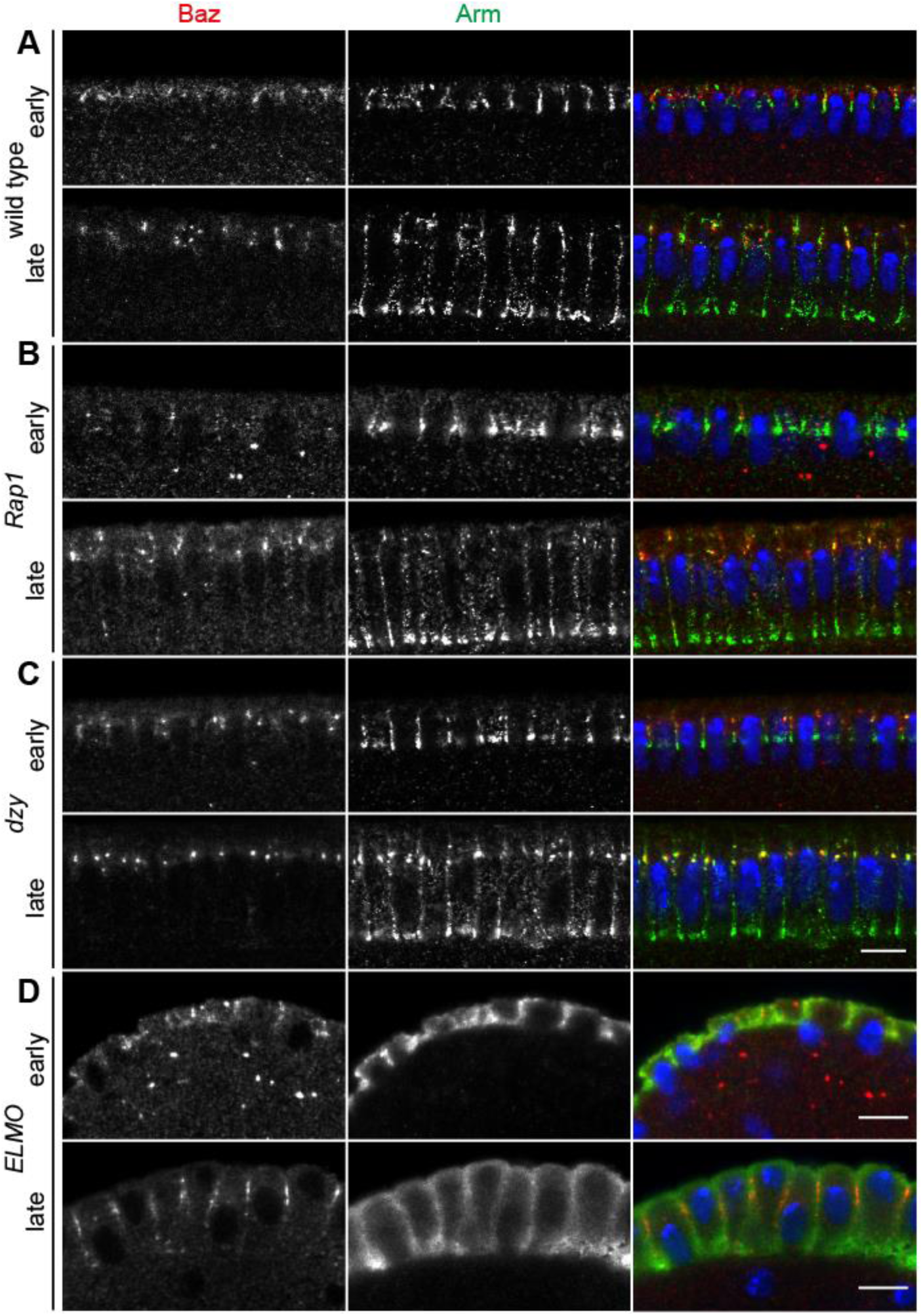
Subapical enrichment of Baz and Arm is perturbed in *Rap1* and *ELMO* but not in *dzy* mutants. (A–D) Fixed cellularizing wild type (A), *Rap1* (B), *dzy* (C) and *ELMO* (D) embryos stained against Baz (grey/ red), Arm (grey/ green). DNA is shown in blue. (A) Wild type embryos during early and late cellularization showed subapical Baz and Arm enrichment. (B) Baz puncta are spread along the lateral membrane in early and late cellularization of *Rap1* embryos. The subapical Arm enrichment was lost whereas basolateral enrichment was still visible. (C) The subapical enrichment of Baz and Arm is not perturbed in early and late cellularizing *dzy* mutant embryos. (D) Baz and Arm subapical localization is lost in *ELMO* mutants during early and late cellularization. Scale bars 10 µm.

## Discussion

In this study we focused on the function of zygotic gene expression on the formation of the subapical domain during onset of cellularization. Next to the already known fact, that ELMO-Sponge and Canoe localize to newly forming cellularization furrows at onset of interphase 14 (8), we were able to show, that Canoe quickly changes its distribution pattern at “old” cellularization furrows (Figure 7). Although we were able to show Canoe preceding Bazooka to localize to the newly forming furrow and subapical domain, with Bazooka being gradually enriched at the subapical domain during the course of cellularization, we confirmed that Canoe is initially restricted independently of Bazooka. In this analysis we relayed on RNAi mediated depletion, since *bazooka* has an essential early function in germline determination (41–43).

**Figure 7.**
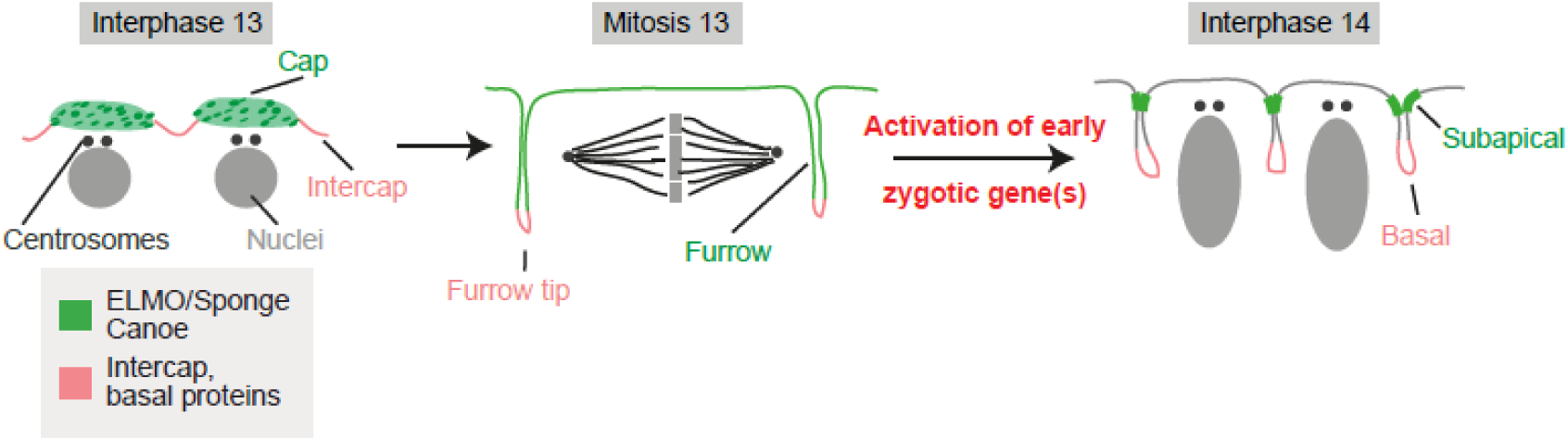
Model of cortical domain protein dynamics from interphase 13 to interphase 14. During interphase 13 cortical domain proteins divide in cap (green) and intercap (red) localization, later subapical proteins like Canoe localize to the whole metaphase furrow during mitosis 13 (green), whereas basal proteins (red) localize to the tip and stay there with remodeling to cellularization furrows. With midblastula transition and onset of zygotic gene expression a yet unknown zygotic gene leads to the remodeling of future subapical proteins in old and new cellularization furrows and subapical domain formation is initiated.

The features of the midblastula transition include deceleration and remodeling of the cell cycle, degradation of maternal products and the switch from syncytial to a cellular blastoderm and the onset of zygotic gene expression (13,36,44,45). As new feature of the morphological changes associated with the midblastula transition we describe here a change in cortical patterning, i. e. the emergence of the subapical domain. Although the restriction of subapical markers precedes formation of a morphologically visible furrow, the apposition of two plasma membranes in initial furrow formation could be the cause of marker restriction, given the limited morphological resolution of our assays. A hint could come from the “old” cellularization furrows that arise from metaphase furrows, which were still detectable in α-amanitin injected embryos by ELMO-GFP. Even at the positions of the old furrow the spatial restriction is lost. A limitation to this argument is again the limited insight into the cellular morphology and dynamics, as the dynamics of the metaphase furrow in embryos lacking zygotic transcription is not clear. A more defined insight into the timing by zygotic gene expression comes from our investigations of embryos with precocious onset of zygotic gene expression. We could detect Canoe at forming cellularization furrows whenever zygotic gene expression was initiated.

The next arising question is which zygotic gene or genes could be responsible for relocalization of the subapical domain proteins with onset of cellularization. Among the described early zygotic genes like *slam, nullo, bottleneck and serendipity- α* no such phenotypes have been described yet (8,18,46–48). However, as general morphology was the primary assay for the screen of zygotic genes (49,50), the subapical determinant might have been missed. A molecular screen of aneuploid embryos for mislocalization of subapical domain proteins may allow the identification of these genes, for example. Although *bazooka* is already maternally expressed, it seems to take over the function as the subapical determinant only later during in cellularization (15,16). Although, it is not clear how much the expression levels were reduced in *bazookaRNAi* embryos, we were not able to detect Bazooka protein by staining.

Taken together, we were able to show, that the formation of the newly established subapical domain is a novel feature of the midblastula transition, which depends on the onset of zygotic transcription. We propose the hypothesis, that a yet unknown zygotic gene triggers the signaling cascade for subapical domain formation involving ELMO-Sponge, Rap1 and Canoe.

## Materials and Methods

### Fly stocks and handling

Fly stocks used were CanoeYFP (*cno[CPTI000590]*, Drosophila Genomics and Genetic Resources, Kyoto), UASp-CherrySlam driven by maternal Gal4 (18), *rap1[P5709]* (R. Reuter, University of Tübingen, Germany)(19), *dizzy(Δ8)* (R. Reuter) (20), *RPII215[X161]* (9), *sésame* (*Hira[185b]*) (21); UASbazRNAi (Bloomington stock # 35002), maternal triple driver MTD-Gal4 (22). As wild type control *w[1118*] was used.

All fly stocks were provided by the Drosophila Stock Center, Bloomington, if not stated differently. Genetic markers and annotations are described in Flybase (http://flybase.org)(23). All crosses and cages were kept at 25°C. Germ line clones were produced with the *ovo*/Flipase technique as described previously (24). *Bazooka* was depleted by overexpression of a short hairpin RNA with MTD-Gal4 during oogenesis.

### Immunostainings and antibodies

Following primary antibodies were employed: mouse anti-Armadillo (1:50; N27A1, Hybridoma Center); rabbit anti-Bazooka (1:1000; A. Wodarz)(25); rabbit anti-Canoe (1:1000; (15)); mouse anti-Dlg (1:100; 4F3, Hybridoma Center); guinea pig anti-Sponge (1:1000; (26)). F-actin was stained by Phalloidin coupled to Alexa647 (Thermo Fisher). Secondary antibodies were labeled with Alexa 488, 568, 647 (Thermo Fisher). GFP tagged proteins was detected with GFP-booster coupled with Atto488 (1:500; Chromotek). DNA was stained by DAPI (0.2 µg/ml; Thermo Fisher).

Embryos were fixed by 4% formaldehyde or by heat fixation using standard methods described previously (27) and stored in methanol at –20°C. For F-actin staining with phalloidin and in the α-amanitin experiments, embryos were fixed by 8% formaldehyde and manually released from the vitelline membrane. For staining, embryos were transferred to PBT (Phosphate buffered saline (PBS) + 0.2% Tween20), washed trice for 5 min and afterwards blocked for 30–60 min in PBT+5% bovine serum (BSA). Embryos were incubated with primary antibodies in PBT+0.1% BSA overnight at 4°C or for 2–3 h at room temperature. Afterwards the embryos were washed with PBT trice for 15 min, incubated with secondary antibodies in PBT for 1– 2 h at room temperature and again washed 3× with PBT for 15 min and stained with DAPI for 10 min at room temperature. The embryos were mounted in Aquapolymount (Thermo Fisher).

Injection of α-amanitin for inhibition of RNA polymerase II was conducted with a concentration of 1 mg/ml in water according to standard procedures as described before (28,29). Afterwards, the embryos were staged to reach interphase 14/15 and fixed as described above. The vitelline membrane was manually removed prior to the staining procedure.

### Imaging and Software

Imaging was performed with a Zeiss LSM780 confocal microscope equipped with an Airyscan detector unit. Fixed samples were imaged with an LCI Plan Neofluar 63×/water NA 1.3 objective. Live imaging was conducted with a Plan Neofluar 63×/oil NA 1.4 objective. Embryos for live imaging were prepared as previously described (30). Fixed samples were imaged with a frame size of 512×512 pixel (67.5×67.5 µm; 130 nm lateral pixel size) for top views and 512×200 pixel (96.4×29.4 µm; 190 nm lateral pixel size) for side views. Top views were conducted as z-stacks with a step size of 0.5 µm. Live imaging was conducted in the Airyscan mode with a frame size of 376×376 pixel (31.7×31.7 µm, 80 nm lateral pixel size). Top views were conducted as axial stacks with a step size of 0.25 µm. Orthogonal views were constructed in Fiji/ImageJ (31). Image were processed in Fiji/ ImageJ, Adobe Photoshop and Illustrator.

## Acknowledgement

We are grateful to E. Geisbrecht, R. Reuter, M. Peifer and A. Wodarz for materials or discussions. We acknowledge service support from the Developmental Studies Hybridoma Bank created by NICHD of the NIH/USA and maintained by the University of Iowa, the Bloomington Drosophila Stock Center (supported by NIH P40OD018537). This work was in part supported by the Deutsche Forschungsgemeinschaft (DFG SPP1464 GR1945/4-2 and equipment grant INST1525/16-1 FUGG).

## Supplemental figures

**Supplemental figure 1.**
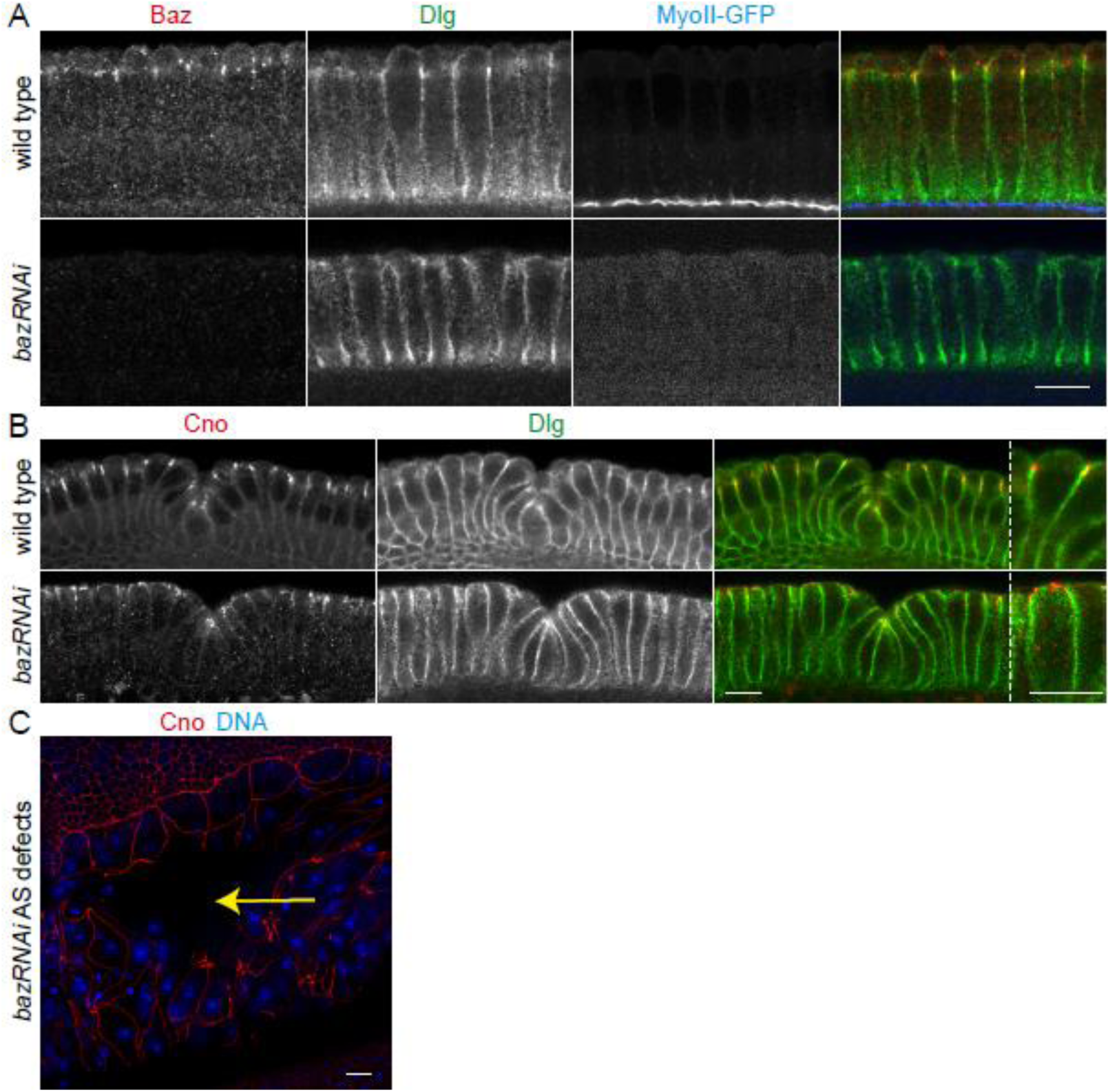
Functionality of *bazooka* knock down. (A) Wild typic embryos expressing MyoII-GFP and *bazRNAi* embryos fixed and stained during late cellularization agains Baz (grey/red), Dlg (grey/ green) and MyoGFP (grey/ blue). MyoGFP and bazRNAi embryos were fixed and stained in the same tubes and imaged with same settings. (B) Fixed wild type and bazRNAi embryos during early gastrulation stained against Cno (grey/ red) and Dlg (grey/ green). (C) Stage 13 bazRNAi embryo fixed and stained against Cno (red) and DNA (blue) showing typical amnioserosa holes (yellow arrow). Scale bars 10 µm.

